# Imaging of single mRNA translation repression reveals diverse interactions with mRNP granules

**DOI:** 10.1101/332692

**Authors:** Stephanie L. Moon, Tatsuya Morisaki, Anthony Khong, Kenneth Lyon, Roy Parker, Timothy J. Stasevich

## Abstract

During cellular stress mRNAs exit translation and accumulate in stress granules and P-bodies, although the dynamics of these interactions remain unclear. We imaged in real-time single mRNAs, their translational output, and mRNA-granule interactions during stress. We observed single mRNAs interacting with stress granules and P-bodies, with mRNAs moving bidirectionality between them. While translating mRNAs only interact with RNP granules dynamically, non-translating mRNAs can form stable associations that rigidly immobilize the mRNA within the granule. Imaging thousands of individual mRNA-granule interactions showed the probability of stable association increases with both mRNA length and granule size. Therefore, the recruitment of mRNAs to RNP granules involves both highly dynamic and stable interactions, influenced by several parameters, demonstrating a new layer of complexity in mRNA regulation during stress.

**One Sentence Summary:** mRNAs interact with stress granules and P-bodies in stable and dynamic manners influenced by ribosome association, mRNA length, and granule size.

## Main text

To survive environmental stress, cells reduce bulk protein production to promote translation of stress responsive proteins. During this response, mRNAs accumulate into cytoplasmic granules of non-translating mRNAs and proteins (RNPs), referred to as stress granules (SGs) and P-bodies (PBs) (*1*). While it is known that mRNAs within these mRNP granules are translationally repressed, is it not known when or where translation is repressed, how dynamic mRNA-granule interactions are, and the order of mRNA association with PB and SG. Resolving these issues are important because mRNA-granule interactions are implicated in multiple processes beyond the modulation of the stress response (*2*, 3), including maternal mRNA storage in early development (*4*), synaptic plasticity (*5*), tumor progression (*6*, 7), and neurodegeneration (*8*–10).

To quantify the relationship between translation of single mRNAs and their interactions with mRNP granules in living cells, we used Nascent Chain Tracking (*11*). The translation of individual mRNAs labeled by fluorescent MS2 coat protein (MCP-Halo with JF646 dye (*12*)) was monitored via the binding of fluorescent antibody fragments (Fab conjugated with the Cy3 dye) to epitopes at the N-terminus of the nascent peptide (Fig. 1A). Thus, translating mRNAs were labeled by both MCP and Fab, while non-translating mRNAs were only labeled by MCP. By imaging these constructs in U-2 OS cells stably expressing the SG marker GFP-G3BP1 (*13*), we examined the translation status of single mRNAs during arsenite stress in relation to their interactions with SGs.

**Fig. 1.**
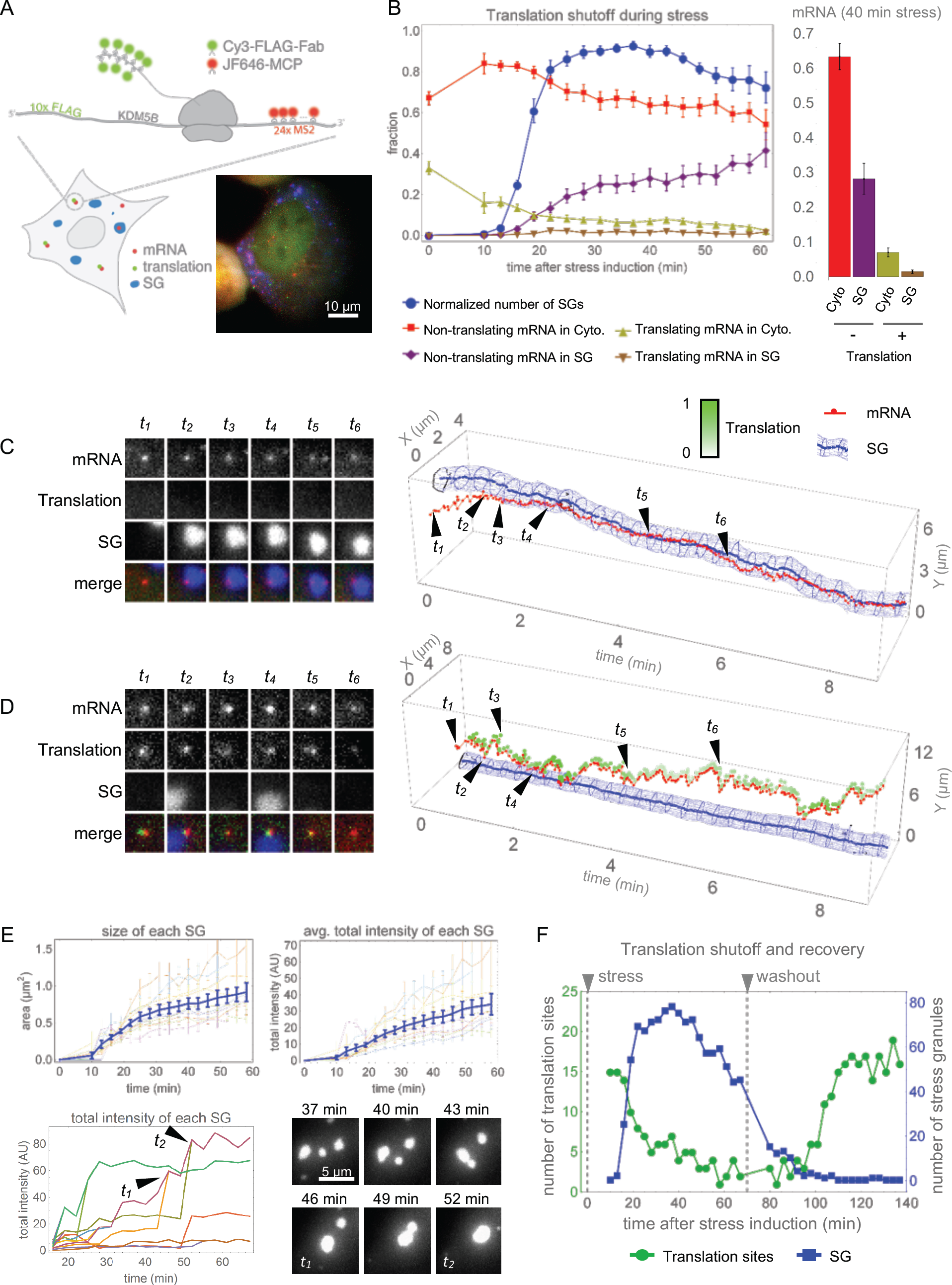
mRNAs cease translation prior to entry into SGs and resume translation following SG disassembly. (A) Single mRNAs with 24X MS2 stem loops in the 3’UTR are visualized with JF646-MCP (red) and translation observed by a-FLAG Cy3-Fab (green) binding the N-terminal ORF 10x FLAG tags in U-2 OS cells expressing the SG marker GFP-G3BP1 (blue); a representative cell is shown. (B) Simultaneous detection of mRNA localization, translation activity, and SG formation in arsenite-stressed cells. Left, normalized number SG per cell (blue); fraction cytoplasmic non-translating (red) and translating (yellow) mRNAs; fraction SGlocalized translating (brown) and non-translating (purple) mRNAs. Right, fraction cytoplasmic or SG localized translating or non-translating mRNAs 40 minutes post-stress. Avg. ±SEM shown from 10 cells. (C and D) Left, representative images of single mRNAs (red), their translation activity (green), and SG (blue) interactions. Right, graphical representation of interactions with intensity of translation foci represented as white-green. (C) A long-term mRNA-SG interaction (>10 min). (D) An example of transient translating mRNA-SG interactions. (E) SG growth over time, as SG average size (upper left) and average SG intensity (upper right). Growth of individual SGs (lower left) with fusion events shown graphically (*t*_*1*_ and *t*_*2*_) and as representative time series (lower right). Avg. ±SEM shown from 10 cells. (F) Representative cell showing translation resumption after arsenite washout. Number of translation foci (green) and SGs (blue) during stress (0-70 min) and following washout (80-140 min).

We first imaged cells at lower temporal resolution (1 volume every 3 minutes for 1 hour) to determine the timing of translation repression during stress. The translation of single KDM5B reporter mRNAs declined upon stress, with SGs forming after 10 minutes of stress (Fig. 1B, Movie S1). The interaction of KDM5B mRNAs with SGs slightly lagged behind SG assembly, reaching a maximum of ∼30% of the mRNAs associating with SGs by 40 minutes of stress, which is similar to what is predicted of endogenous KDM5B mRNAs from the SG transcriptome (∼39%) (*14*). 98% of the KDM5B SG-associated mRNAs were not translating, implying translation repression is a general prerequisite for mRNA recruitment to SGs. We observed 1-2% of SG-associated mRNAs retained nascent chains, suggesting that mRNAs can interact with SGs while still associated with ribosomes. To verify these observations, we repeated experiments at a higher temporal resolution (1 volume every 2 seconds for 10 minutes) between 10 and 30 minutes post-stress, a period when both translating mRNAs and SGs are observed (Fig. 1B). This confirmed that most of the mRNAs entered SGs in a translationally repressed state (Fig. 1C, Movie S2). On rare occasions, translating mRNAs (with nascent chains) interacted with SGs transiently (Fig. 1D; Movie S3). Most of these transient interactions were on the order of seconds, but a small subset could last up to a few minutes (Fig. S1, Movie S3).

The repression of mRNA translation was positively correlated with the growth of SGs, which increases throughout the stress response (*15*–17). Evaluation of SGs revealed SG growth by both incremental accumulation of material and SG fusion (Fig. 1E). Incremental growth dominated during the early phases of stress, whereas fusion events dominated at later times. SG growth also correlated with increased KDM5B mRNAs in SGs, consistent with mRNA recruitment contributing to SG growth.

We then examined the kinetics of translation resumption during the recovery from arsenite stress. Stress granule disassembly (quantified as number of individual SGs) occurred just a few minutes prior to translation resumption in single cells (Fig. 1F, Fig. S2, Movie S4). We verified that these translating spots in the recovery phase were bona-fide translating mRNAs by observing their disappearance following puromycin treatment (Movie S5). While KDM5B mRNAs ceased translation gradually as SGs formed, during the recovery phase KDM5B did not resume translation until stress granules had dissolved completely. These observations suggest that SGs disassemble prior to translation resumption during stress recovery.

We next investigated the dynamics of mRNA-SG interactions by measuring the duration of individual SG interactions with mRNAs encoding either KDM5B or H2B (Fig. 2A&B). In both cases, the probability an SG-mRNA interaction lasted a set duration of time (i.e. the interaction time survival probability) could be fit by a two-state model with fast interactions on the order of seconds and slow interactions on the order of many minutes (Fig. 2C&D). These results suggest mRNAs frequently ‘sample’ SGs and occasionally enter a stable association.

**Fig. 2.**
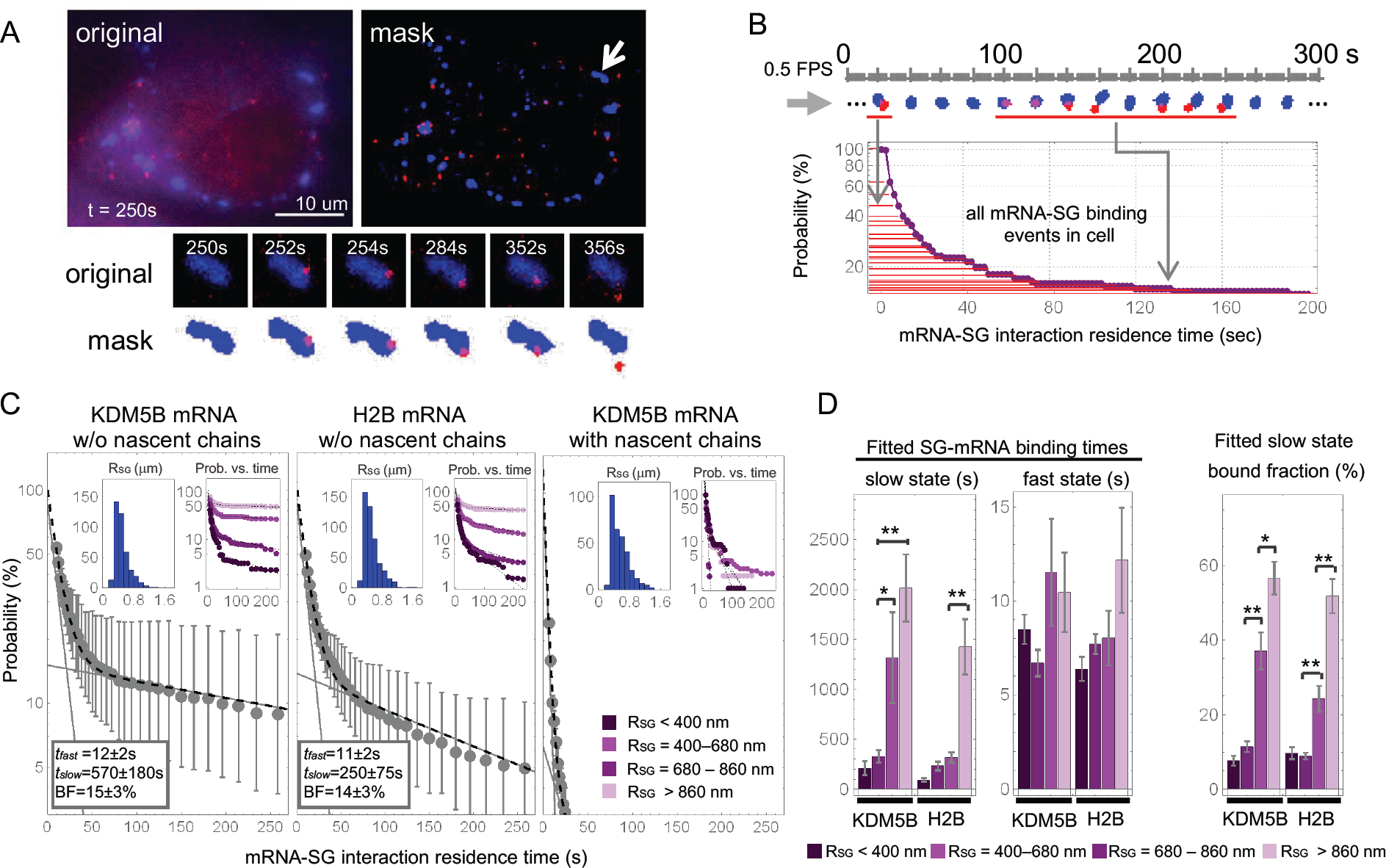
mRNAs interact transiently and stably with SGs. (A) Determination of mRNA-SG interaction times. Representative image showing mRNAs (red) and SGs (blue) (upper left) and the masked image (upper right) with an example mRNA-SG interaction (arrow) below. (B) Survival probability distribution of mRNA-SG interaction times (red) from a representative cell. (C) The survival probability of non-translating KDM5B (left; 9 cells, 409 tracked SGs) and H2B (middle; 11 cells, 492 tracked SGs) mRNA-SG interactions has a fast and slow component, while translating KDM5B mRNA-SG interactions has a fast component (right; 9 cells, 326 tracked SGs). Inset histograms depict the distribution of stress granule radii *R_SG_* from the masked image (μm) and inset graphs depict the survival probability distribution of mRNA-SG interaction times for a given effective SG radius (right panel legend). Avg. fitted fast and slow interaction times and slow fractions summarized in the left bottom insets (±90% CI). (D) Avg. fitted slow & fast interaction times, and slow fractions for a given effective SG radius (±90% CI). * p< 0.01; ** p< 0.001.

More generally, three parameters influenced the nature of SG-mRNA interactions. First, we observed that the H2B reporter mRNA, whose ORF is ∼1400 codons shorter than the KDM5B ORF, interacted with SGs the same as KDM5B in the fast state, but was not as stably associated with SGs in the slow state (Fig. 2C, t_fast_ = 12 ± 2 s vs. 11 ± 2 s and t_slow_ = 570 ± 180 s vs. 250 ± 75 s). We also observed that the duration of the slow interactions and the fraction of slow interactions for both the KDM5B and H2B mRNAs varied depending on the size of the SG. Larger SGs showed more and longer stable interactions with mRNAs than smaller SGs (Fig. 2C&D, Movie S6 & S7). Finally, we observed that mRNAs still engaged with ribosomes, as assessed by the detection of nascent peptides, could interact with SGs transiently but did not engage in the stable interaction mode (Fig. 2C, right panel).

To determine if other mRNP granules also show bimodal interactions with mRNAs, we examined the dynamics of mRNA interactions with PBs, as marked with mRFP-DCP1a. We observed that both KDM5B and H2B mRNAs showed rapid and stable interactions with PBs (Fig. 3A), that KDM5B showed longer stable interactions than H2B mRNAs (Fig. 3A&B), and that the degree of stable interactions for either mRNA increased with the size of the PB (Fig. 3A&B; Movie S8). These observations indicate that mRNAs can dock and undock to SGs and PBs through transient interactions, and in some cases form a stable set of interactions that ‘locks’ the mRNA into the granule for prolonged periods. Stable association could be more prevalent on longer mRNAs and in larger SGs and PBs due to increased sites for additional interactions. This model implies that the association of an mRNA with a granule can be influenced by *cis* or *trans* inputs that modify either the docking/undocking step, or the rate of entry into a stable, locked interaction state, which is supported by the observation that knockdown of the RNA binding protein LARP1 can reduce the association of some mRNAs with PBs and SGs (*18*).

**Fig. 3.**
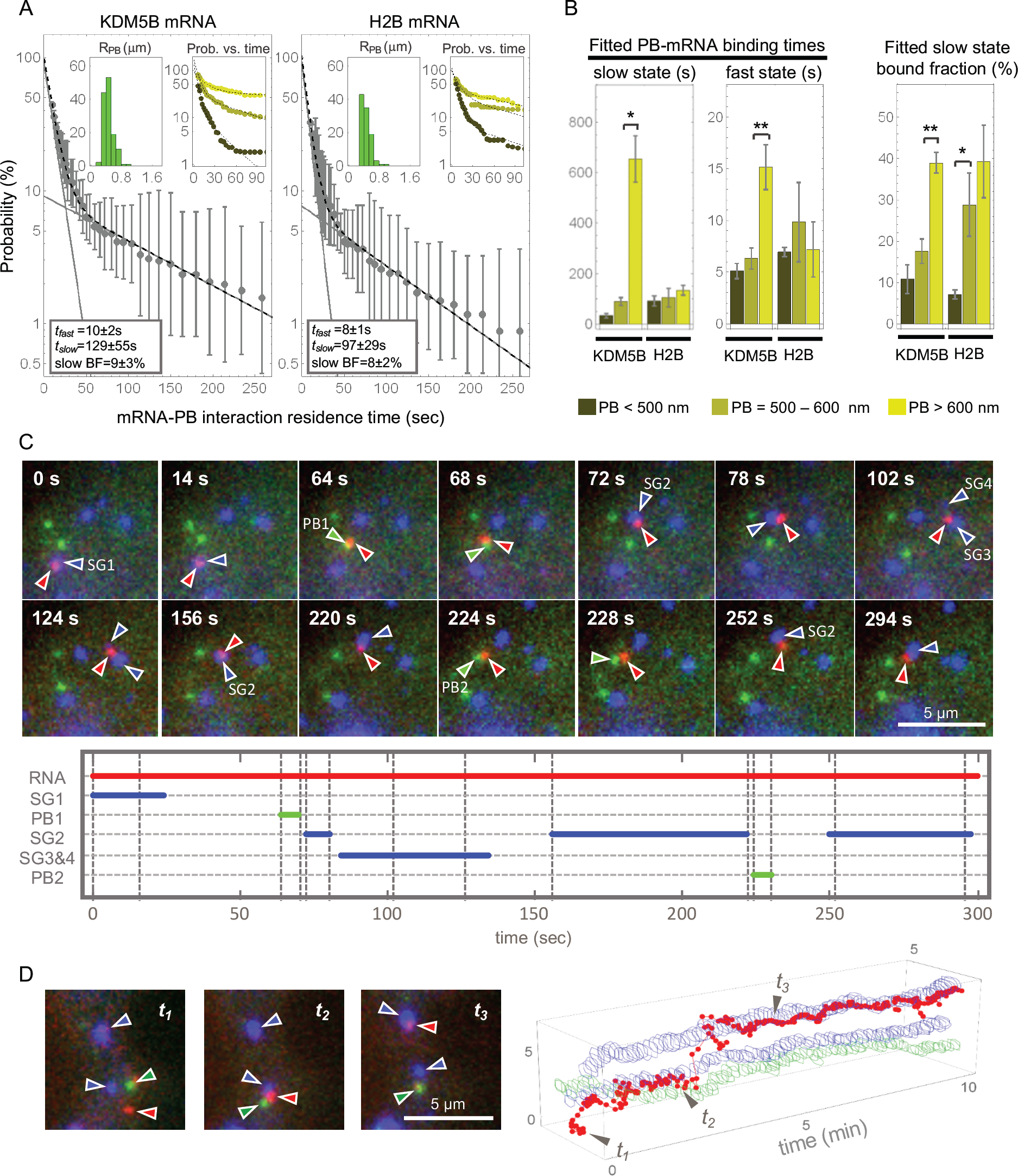
mRNAs interact transiently and stably with PBs and traffic bidirectionally between PBs and SGs. A) Avg. survival probability distributions of KDM5B mRNA-PB (left; 7 cells, 137 tracked PBs), and H2B mRNA-PB (right; 4 cells, 106 tracked PBs) interaction times ±SEM. Inset histograms depict the distribution of effective PB radii from masked images (µm), and inset graphs depict the survival probability distributions of mRNA-PB interaction times for a given effective PB radius (Fig. 3B, bottom). Avg. fitted fast and slow interaction times and slow fractions are in left bottom insets ±90% CI. (B) Avg. fitted slow and fast interaction times and slow fractions for a given effective PB radius ±90% CI. * p≤ 0.01; ** p≤ 0.001. (C) Representative images (top) of a single KDM5B mRNA (red, red arrows) that interacted with four SGs (blue, blue arrows) and two PBs (green, green arrows). The duration of each mRNA-RNP granule interaction was plotted (bottom). (D) An mRNA trajectory between two SGs and a PB visualized by plotting the position of the mRNA (red), SG (blue), and PB (green) over time. Example cropped images corresponding to time points *t*_*1*_ to *t*_*3*_ are shown (left).

Non-translating mRNAs can accumulate in both SGs and PBs. A general model is that following translation repression mRNAs first associate with SGs, and then can be sorted for targeting to PBs (*19*). In contrast, it has been proposed that mRNAs move from PBs to SGs upon glucose deprivation in yeast (*20*). To determine if there is a preferred movement of mRNAs between SGs and PBs, we simultaneously imaged SGs, PBs and KDM5B mRNAs during stress. We observed mRNAs exchange between SGs and PBs in a bidirectional manner (Fig. 3C&D). These interactions included cases of mRNAs transitioning from a SG to a PB and back to a different SG within as little as ∼30 seconds (Fig. 3C, Movie S9) and instances where the mRNA was localized between a PB and a SG for a few minutes (Fig. 3D & Movie S10). These rapid bidirectional movements between SGs and PBs indicate that there is no obligate path for mRNA movement between these granules, which also demonstrates that an individual mRNA with it’s associated proteins is capable of interacting with either SGs or PBs, or exchanges proteins that enable SG or PB interaction within seconds.

SGs, and other mRNP granules, have been proposed to be liquid-like compartments with components showing rapid diffusion within the compartment (*21*–24) and/or contain liquid-like areas surrounding densely compacted ‘cores’ (*15*, 25). Those mRNAs that were stably associated with mRNP granules for many minutes adopted the motility of SGs (Fig. 4A) and could be observed at the surface or embedded within SGs (Fig. 4B). To examine the liquid-like nature of SGs, we examined the movement of mRNAs relative to one another within SGs. Surprisingly, we observed multiple cases where the relative localization of mRNAs, as assessed by quantifying the distance and angle between three individual KDM5B mRNAs in one SG, remained relatively constant over time (Fig. 4B&C, Fig. S3, Fig. S4, Movie S11), consistent with limited movement of the mRNA within the larger assembly. Quantification revealed the intra-SG diffusion of mRNAs to be 280 times slower than the SG as a whole and over 1600 times slower than free mRNAs within the cytoplasm (Fig. 4A). These observations demonstrate that mRNAs within large SGs can be rigidly positioned, suggesting they are within a non-liquid-like environment.

**Fig. 4.**
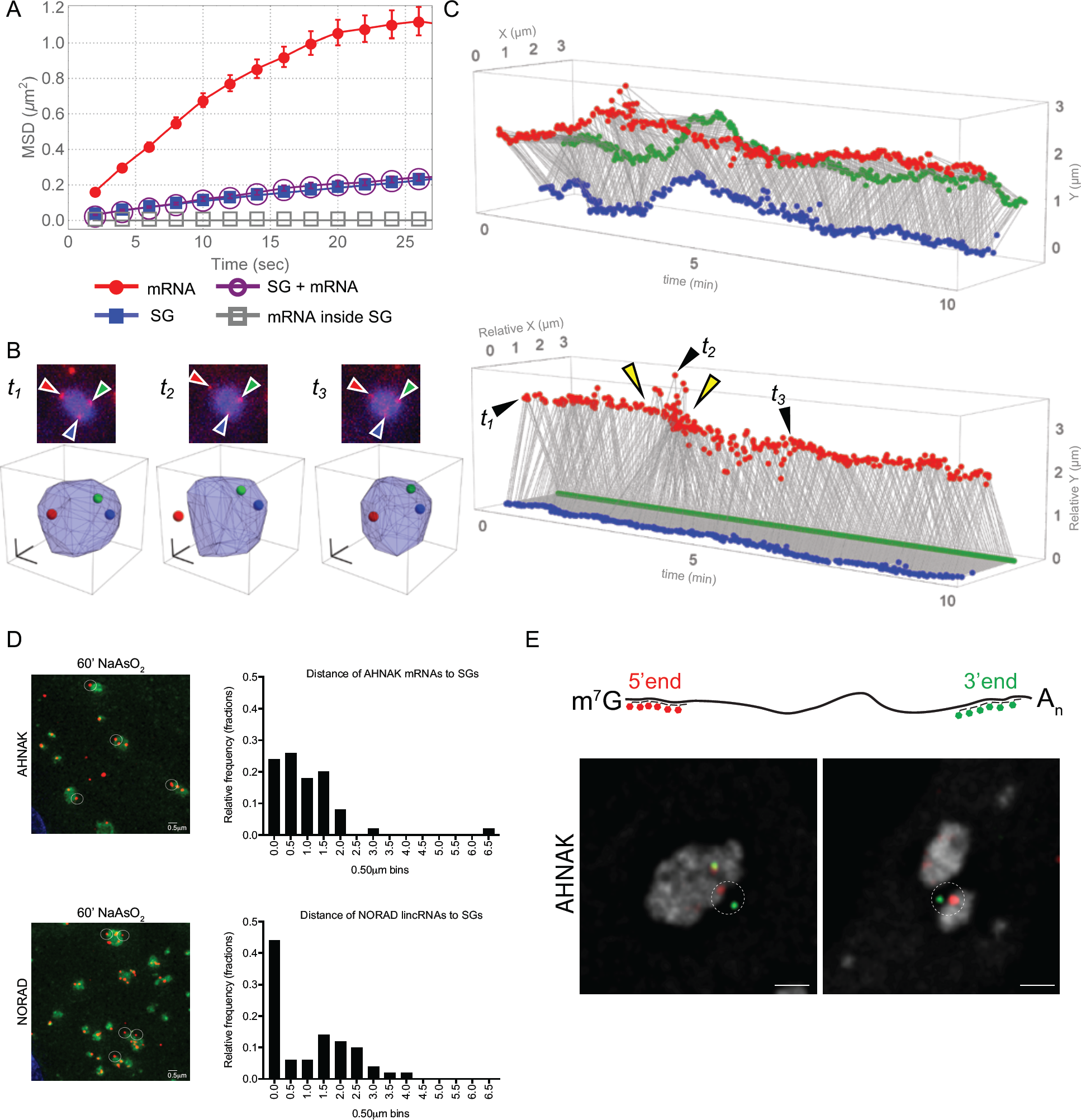
mRNAs can be rigidly positioned within SGs and/or tethered to them. (A) Average mean squared displacement (MSD) of cytoplasmic KDM5B mRNAs (red; fitted D = 0.016 ± 0.001 µm^2^/sec), SGs (blue; fitted D = 0.0024 ± 0.0003 µm^2^/sec), SGs containing mRNAs (purple circles; fitted D = 0.0028 ± 0.0004 µm^2^/sec), and mRNAs in SGs (gray squares; fitted D = 10 ± 2 nm^2^/sec) from a representative cell (163 tracked mRNAs, 250 tracked SGs, 13 tracked SGs containing mRNAs). (B) A representative image time series showing three mRNAs (red foci marked by red, blue and green arrows) in a SG (top) and their 3D positions (as red, green or blue dots) inside the SG (bottom). Scale bars 1 µm. (C) Projected 2D positions of the mRNAs depicted in Fig. 4B plotted over time (top). Relative 3D positions of the three mRNAs plotted over time by fixing the position of the green mRNA and the orientation of the blue mRNA with respect to the green mRNA (bottom). Yellow arrows indicate exit and entrance of the red-labeled mRNA from the SG. D) Sample immunofluorescence-smFISH images acquired for AHNAK and NORAD mRNAs (red) and SGs (G3BP1, green) at 60 min. post stress. Left, representative images of mRNAs clustered near SGs (white circles). Scale bar, 0.5µm. Right, histograms depict relative frequency of measured distances of 50 individual AHNAK or NORAD RNAs to nearest SGs. E) Cartoon of positions of smFISH probes along AHNAK mRNA (top). (Bottom) Representative smFISH images where one end of the AHNAK mRNA was outside the SG while the other end was inside the SG (5’ end probes in red, 3’ end probes in green; SG in grey). Scale bars, 1µm.

Unexpectedly, we observed multiple cases where the MCP signal, which represents the 3’ end of a KDM5B mRNA, exited the SG and then returned to approximately the same position within the SG (Fig. 4C, yellow arrows, Movie S11, Fig. S3, Fig. S4). This observation suggested that the 3’ end of this mRNA might exit the GFP-G3BP1-marked portion of the SG while the remainder of the mRNA is still within the SG. Consistent with the body of an mRNA extending beyond the G3BP1-marked SG, we observed that endogenous RNAs that localize to SGs (AHNAK and NORAD) are often detected near, but not in SGs in U-2 OS cells by smFISH (Fig. 4D), with the shorter NORAD RNA (∼5 kb) being closer to the periphery of a SG than the longer AHNAK RNA (18kb). Furthermore, using smFISH probes to the 5’ or 3’ ends of the AHNAK mRNA, we observed multiple instances where one end of the mRNA was outside of the SG while the other end co-localized with GFP-G3BP1 inside the SG (Fig. 4E). These observations suggest that SG-associated mRNAs can extend beyond SG boundaries as defined by protein components, and thus could provide a set of interactions to promote fusion of smaller SG and docking of SG and PBs, which is consistent with our observations of individual mRNAs localizing between a PB and a SG for several minutes (Fig. 3D & Movie S10).

These experiments demonstrate that mRNAs interact in both stable and transient manners with SG and PBs. In model assemblies, exchange rates of components are related to both the strength of individual interactions, and the valency, with higher valency having more stable interactions (*26*). This suggests that the transient interactions with SG or PB represent a low valency mode of interaction but provide a “docked” state whereby mRNPs can form additional interactions with SG components to enter a stable “locked” state with higher valency (Fig. S5). Higher valency would be expected on longer mRNAs, and larger SG or PB, providing an explanation for why longer mRNAs and larger RNP granules have more stable RNA-granule interactions. Our experiments provided unambiguous data that mRNAs stably associated with SG are not translating, although mRNAs associated with ribosomes can transiently interact with SG. This may be because the 3’UTR and/or partially exposed coding regions may be sufficient to form transient protein-protein or RNA-RNA interactions, but a fully exposed coding region is required to make sufficient interactions for stable SG association. We suggest that this bimodal nature of RNA-granule interaction will be a general principle of any RNP granule where the RNA enters a multivalent state. This work identifies the docking and locking steps as two distinct steps in RNP association with any RNP granule that can be modulated to affect RNA-granule association.

## Acknowledgements

We thank Nancy Kedersha for providing GFP-G3BP1 mRFP-DCP1a U-2 OS cells and Esther Braselmann and Theresa Nahreini for isolating GFP-G3BP1 U-2 OS cells at the BioFrontiers Institute Flow Cytometry Core facility. We thank lab members for helpful suggestions.

## Author Contributions

Conceptualization, S.L.M., R.P.; Methodology, S.L.M., T.M., T.J.S., A.K., K.L.; Software, T.M., T.J.S.; Validation, S.L.M.; Formal Analysis, T.M., T.J.S., S.L.M., A.K.; Investigation, S.L.M., A.K., K.L., T.M., T.J.S.; Resources, R.P., T.J.S., T.M., A.K., K.L., S.L.M.; Data Curation, T.M., T.J.S., S.L.M.; Writing-Original Draft, S.L.M., R.P.; Writing-Review & Editing, R.P., S.L.M., T.J.S., T.M., A.K.; Visualization, T.M., T.J.S., S.L.M., A.K.; Supervision, R.P., T.J.S., T.M., S.L.M.; Project Administration, S.L.M.; Funding Acquisition, T.J.S., R.P., S.L.M.

## Data and materials availability

All data are included in the manuscript and/or available upon request.

## Supplementary Materials

Materials and Methods

Figure S1-S5

Movies S1-S11

